# Coarse, Medium or Fine? A Quantum Mechanics Approach to Single Species Population Dynamics

**DOI:** 10.1101/2020.07.22.215061

**Authors:** Olcay Akman, Leon Arriola, Aditi Ghosh, Ryan Schroeder

## Abstract

Standard heuristic mathematical models of population dynamics are often constructed using ordinary differential equations (ODEs). These deterministic models yield pre-dictable results which allow researchers to make informed recommendations on public policy. A common immigration, natural death, and fission ODE model is derived from a quantum mechanics view. This macroscopic ODE predicts that there is only one stable equilibrium point 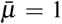. We therefore presume that as *t* → ∞, the expected value should be 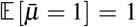. The quantum framework presented here yields the same standard ODE model, however with very unexpected quantum results, namely 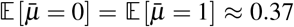. The obvious questions are: why isn’t 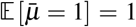, why are the probabilities ≈ 0.37, and where is the missing probability of 0.26? The answer lies in quantum tunneling of probabilities. The goal of this paper is to study these tunneling effects that give specific predictions of the uncertainty in the population at the macroscopic level. These quantum effects open the possibility of searching for “black–swan” events. In other words, using the more sophisticated quantum approach, we may be able to make quantitative statements about rare events that have significant ramifications to the dynamical system.

## 1 Motivation and introduction

Standard temporal population models describe how population changes given the current status of the population. Environmental conditions such as limited resources, competition, disease, etc. affect changes in the population. Additionally, processes such as birth, death, immigration, and emigration affect changes to the age–size–distribution. These standard models are based on heuristic arguments. In other words, the models are based on the modeler carefully deciding which are the most important aspects of the system. Usually population balance equations are heuristically constructed in order to characterize those mechanisms/interactions that the modeler ‘‘thinks” are important to the model.

In 1838 Pierre–Francois Verhulst published a *Note on the law of population growth* [8].

> “We know that the famous Malthus showed the principle that the human population tends to grow in a geometric progression so as to double after a certain period of time, for example every twenty five years. This proposition is beyond dispute if abstraction is made of the increasing difficulty to find food…
>
> The virtual increase of the population is therefore limited by the size and the fertility of the country. As a result the population gets closer and closer to a steady state.”

Verhulst proposed the commonly accepted macroscopic ordinary differential equation

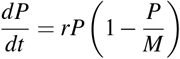

with specified initial condition *P*(0) = *P*_0_ and is commonly referred to as the logistic model. The basic assumption is that the population, *P*(*t*) at time *t*, is homogeneous and uniformly mixed. That is, interactions between any member of the population are equally likely. The heuristic assumptions are the processes of intrinsic birth at rate *r* and limited resources via the carrying capacity *M* respectively.

In this paper, we also use the heuristic approach. However, the essential difference is that the mechanisms and interactions are formulated at the quantum level. For illustrative purposes, we will construct a quantum toy model where we assume the only important quantum interaction is competition as shown in Figure 1.

**Figure 1:**
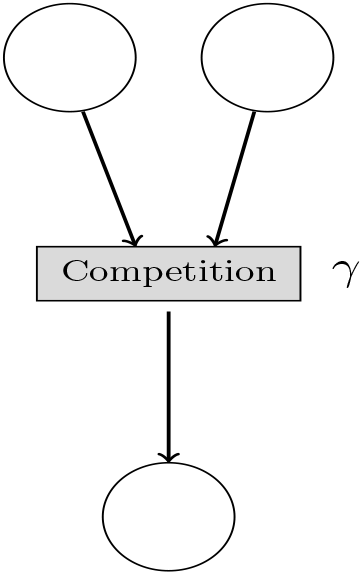
Competition With Rate *γ*

We will define an appropriate Schrödinger equation for which we will find that the time dependent expected value *μ*(*t*) of the “wave function” is governed by the logistic model

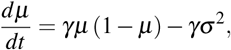

where *σ*^2^ denotes the variance of an underlying probability distribution of the population. Notice the quantum approach yields the familiar macroscopic logistic model with a time varying ‘‘harvesting” term *γσ*^2^ that appears as noise. This effect initially seems to have no relation to assuming competition as the only mechanism of interaction within the population. The quantum approach suggests that the inevitable fluctuations of interactions within the population needs to be included in the standard heuristic macroscopic ODE models. It is also surprising that assuming only competition, the quantum approach yields a growth term *γμ*. From a macroscopic viewpoint, if the only interactions are decay processes, we would not expect growth to occur.

One major goal of this study is to establish an intimate connection between the principals of quantum/stochastic mechanics [2] and the foundations of single species population dynamics. Additionally, we provide a formal framework for a deeper understanding of the underlying processes in single species population dynamics.

For instance, the noise that is predicted by the quantum approach adds a new feature which is not usually included in standard ODE models. Specifically, we propose that the quantum viewpoint validates, explains, and makes specific predictions about the quantum tunneling of probabilities.

Consider another toy quantum model where we assume that the only important quantum mechanisms are immigration, natural death and fission as shown in Figure 2.

**Figure 2:**
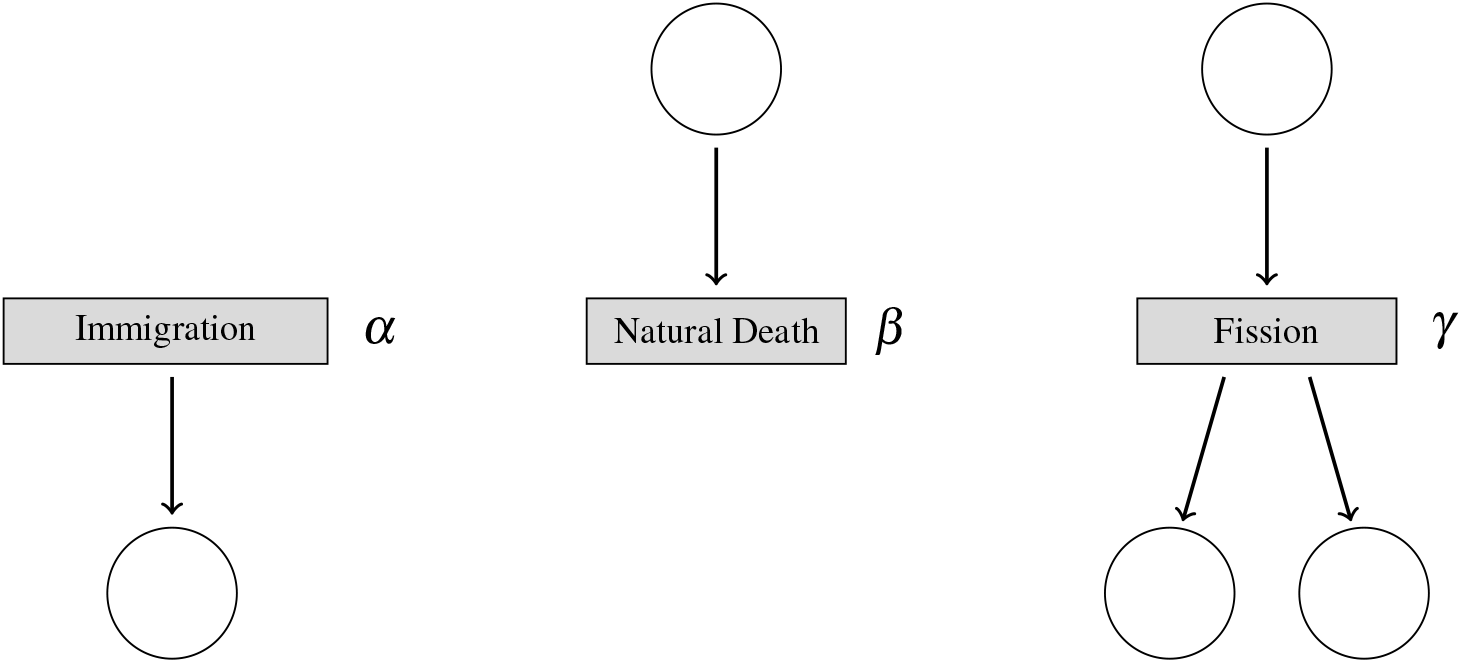
Immigration Birth (*α*), Natural Death (*β*), Fission (*δ*)

The quantum/stochastic mechanics approach will yield the specific Schrödinger equation

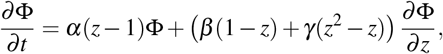

where the Markov generating function **Φ** is defined as

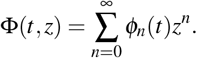

This function describes the temporal probability *ϕ_n_*(*t*) of having exactly *n* objects at any time *t*. The monomials *z^n^* have no physical meaning and can be thought of as place-holders. The expected value of **Φ**, denoted by 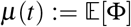, yields the the standard macroscopic linear ODE model

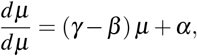

with intrinsic growth rate *γ* – *β* and constant growth *α*. This ODE has the non–negative stable equilibrium 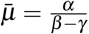, provided *β* > *γ*.

We will examine the very simple case where *α* = *γ* = 1 and *β* = 2, which yields the stable equilibrium *μ* = 1. Let *ϕ*_0_(*t*) denote the probability of having exactly zero objects at time *t*. Similarly, *ϕ*_1_(*t*) denotes the probability of having exactly one object at time *t*, *ϕ*_17_(*t*) denotes the probability of having exactly 17 objects at time *t*, etc.. The standard macroscopic ODE model predicts that *μ*(*t* → ∞) = 1. This suggests that the quantum approach should therefore predict that *ϕ*_1_(*t* → ∞) = 1 and *ϕ*_1_(*t* → ∞) = 0 for all *j* ≠ 1. The quantum approach predicts the very surprising and non–intuitive result

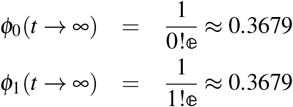

which does not sum to 1! The obvious question is: where is the missing probability 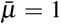 Examining the higher order terms *ϕ_j_*(*t* → ∞) for *j* ≥ 2 we find that

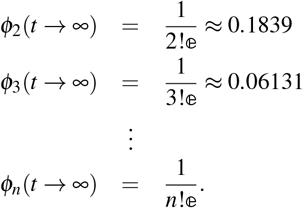

Due to the sophistication of the quantum mechanics paradigm, this means that the probability of having 17 objects is not zero, however it will be very small.

This quantum tunneling effect of probabilities opens the possibility of searching for “black–swan” events. In other words, we may be able to make quantitative statements about rare events that have significant ramifications to the dynamical system. In future work, this framework will be extended to multiple species dynamics such as such as the predator–prey/Lotka–Volterra and the standard susceptible, infected, and recovered (SIR) epidemiological ODE models. The following section contains a short discussion of quantum physics concepts that are applicable to this work.

### 1.1 Quantum physics

One of the most profound paradigm shifts of the 20th century was the quantum theory of physics which was first developed by Bohr, Einstein, Planck, among others [5]. It became quite clear that the elementary processes of physics did not follow the commonly accepted principles of classical mechanics. Prior to the quantum viewpoint, the prevailing notion was that the macroscopic description of nature could be described by an averaging process. As the results of key experiments such as the Youngs’ double–slit experiment were analyzed, the accepted macroscopic viewpoint was shown to be completely inadequate in describing submicroscopic processes.

Most physical systems consist of an astronomical number of individual components along with a corresponding overwhelming number of interactions between the components. Many of the properties of the components often can have significant variation about nominal values. If the modeler was to include all the interactions, a discrete agent model would result which we refer to as a fine scale model. The astronomical number of interactions could not be determined analytically, in which case the modeler would have to resort to computer simulation.

In chemistry, mathematical biology, mathematical epidemiology, population dynamics and other related fields, master equations such as the logistics equation and predator–prey models such as the Lotka–Volterra ODEs, are constructed and used to describe the macroscopic behavior of the time evolution of the dynamical system and will be referred to as a coarse scale model.

These master equations are constructed by making simplified heuristic assumptions in order to define tractable deterministic models. However, these simplifications preclude prediction as well as a deeper understanding about what can occur at the quantum level. In this work we examine in depth the quantum processes as well as the subsequent predictions made to the macroscopic realm–hence a medium scale model.

In the following section we present an a novel approach to address this problem by borrowing concepts of quantum physics.

### 1.2 Feynman diagrams

In order to motivate why we propose a quantum/stochastic mechanics approach to modeling processes, consider the Feynman diagram [7] depicting the interaction of an electron with another electron in a perfectly elastic collision as shown in Figure 3.

**Figure 3:**
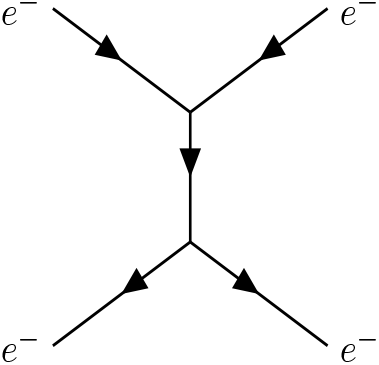
Feynman Diagram of Electron–Electron Interaction

The crucial elements of this diagram are

- objects enter a common location and undergo an interaction with each other and
- objects emerge from the common location after the interaction has occurred.

Consider the four fundamental forces of nature as depicted in the Feynman diagrams as shown in Figure 4.

**Figure 4:**
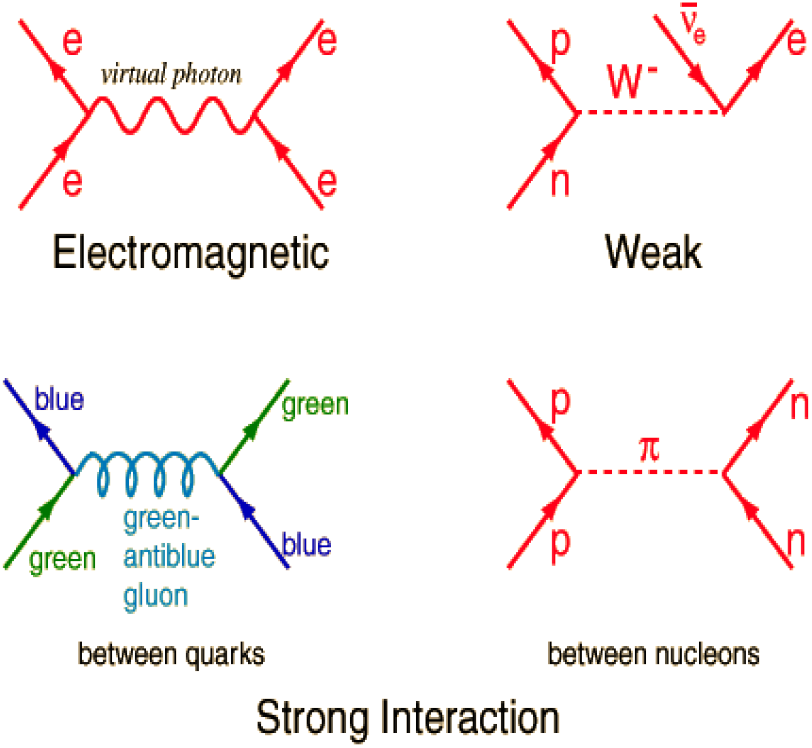
Feynman Diagram of Four Fundamental Forces

Notice that each graph depicts very complex interactions without the burdensome mathematical formalism. Feynman diagrams dramatically validate the old adage: “A picture is worth a thousand words.”

Using the Feynman diagram paradigm, the tools and techniques from quantum physics can be used to describe and understand population dynamics. The beauty of the Feynman diagram approach is that very complex interactions can easily be visualized without the complicated mathematical machinery.

Although the focus of this study is single species population dynamics, it is instructive to point out that future work will provide a natural extension to multiple species interactions such as susceptible or infected populations (SI), or the Lotka–Volterra predator-prey models. For example, consider the interaction between a susceptible and infectious person. Here we assume that the only two possibilities that can occur are the susceptible stays susceptible, or the susceptible becomes infected as seen in Figures 5 and 6, respectively.

**Figure 5:**
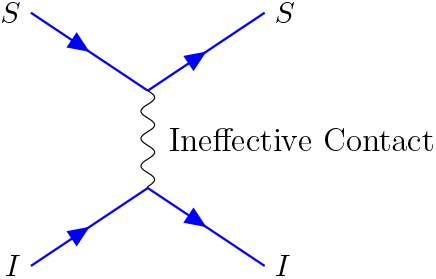
Unsuccessful Transmission of Pathogen

**Figure 6:**
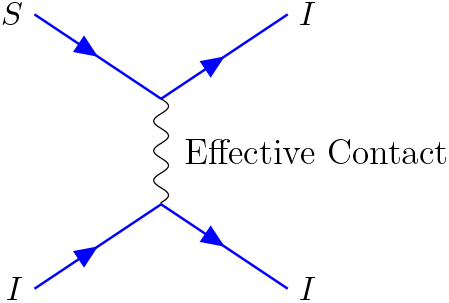
Successful Transmission of Pathogen

### 1.3 Memory–less vs memory processes

Dynamical systems, such as single species populations [1, 3, 4, 6], can evolve whereby the current state of the system has or has no memory of previous states of the system. For example, a Markov chain is an evolution through a sequence of transitions determined entirely by the roll of dice. Ignoring the possibility that the dice have the property of being quantum entangled, previous rolls of the dice cannot possibly affect future outcomes of the dice.

Card games such as blackjack and poker however do have a “memory” of previous states. In fact, a seasoned card player uses this information to their advantage by remembering which cards have been exposed. Deductions can be made by regarding which cards remain in the deck. In the dynamical systems discussed in this paper, the transitions are assumed to be strictly determined by the roll of the dice. In other words, the past, present and future states are statistically independent via a stochastic Markov chain.

### 1.4 Quantum mechanics and population dynamics

In quantum mechanics the actors in the play are the creation operator *a*^+^ and the annihilation^1^ operator *a*^−^. The operator *a*^+^ takes an object from energy level 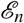 and moves it up one level to 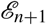. The operator *a*^−^ takes an object from energy level 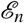 and moves it down one level to 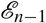. Analogously, in population dynamics we define the creation operator *a*^+^ to take *n* objects and turn them into *n* + 1 objects. The annihilation operator *a*^−^ destroys one of the *n* objects into *n* − 1 objects.

#### 1.4.1 Growth and the creation operator a^+^

Consider the scenario where we initially have three objects *A*, *B* and *C*, and then add one generic external object *D* as seen in Figure 7. Notice there is only one way of adding an additional generic object *D*. Now introduce a monomial *z*^3^ which represents the fact that initially there are exactly three objects. Please note that the independent variable *z* has no physical meaning. The only point of interest is the exponent 3 and the coefficient of the monomial. In other words, we can think of the monomial *z*^3^ as a placeholder. After the interaction has occurred, there are now exactly four objects, which we associate with the monomial *z*^4^. This means that the initial monomial *z*^3^ now becomes *z*^4^. We assume that this holds true for any initial number of *n* objects, in which case by induction the initial monomial *z^n^* becomes *z*^*n*+1^, 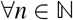. The creation operator *a*^+^ simply multiplies the monomial *z^n^* by *z* yielding *z*^*n*+1^. Hence we define the action of the creation operator *a*^+^ to be

**Figure 7:**
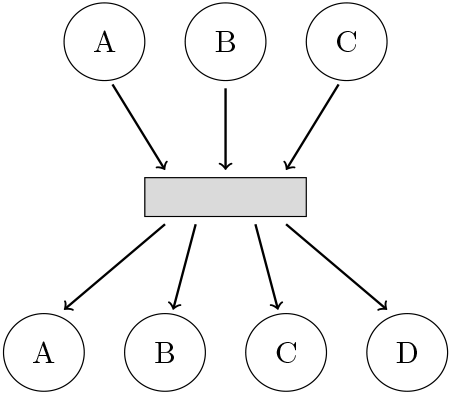
{*A,B,C*} → {*A, B, C, D*}

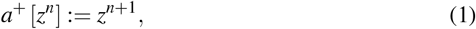

#### 1.4.2 Decay and the annihilation operator *a*^−^

Now consider the reverse scenario where three objects interact, but now one object is annihilated, resulting in two remaining objects. Since there are three ways for two objects to remain in existence, as seen in Figure 8, we associate the action of *a*^−^ to be *a*^−^ [*z*^3^] = 3*z*^2^. We assume this holds true for any non-negative integer number of n objects, the initial monomial *z^n^* becomes *nz*^*n*−1^, in which case the annihilation operator *a*^−^ is just the usual derivative operator *∂*/*∂z*. Hence, we define the action of the annihilation operator *a*^−^ to be

**Figure 8:**
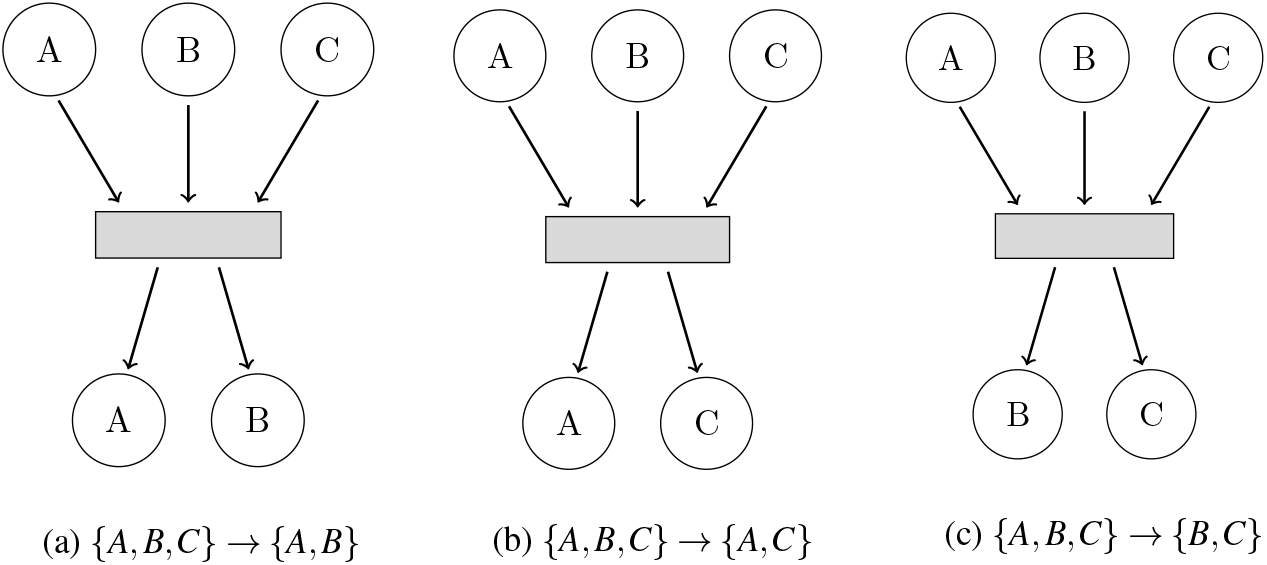
Eliminate One Object From {*A, B, C*}

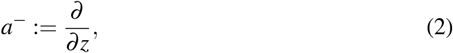

and by induction define

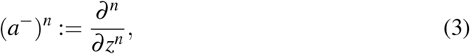

where 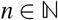. [2].

#### 1.4.3 Combinatorial meaning of the operators a^+^ and a^−^

Consider the interaction where two objects interact sexually and produce a single off-spring with rate *λ* as shown in Figure 9.

**Figure 9:**
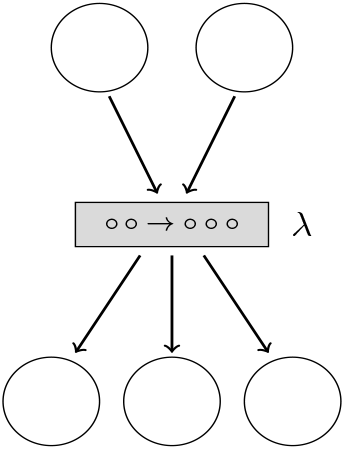
Sexual Reproduction with Rate *λ*

In order to motivate how the number of ways that objects change from the input side *j* = 2 as compared to the output side *k* = 3 we discuss the formulation of the Hamiltonian operator as found in [2]. A Hamiltonian operator can be thought of as a change in energy, a change in probability flux, or in our situation, the change in the number of ways that the input side does not change vs. the number of ways the output changes. To simplify the idea to its most basic form, we casually define the Hamiltonian operator as

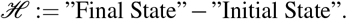

The expressions “Final State” and “Initial State” need to be appropriately defined.

For this example, consider the action of annihilating two objects on the input side, followed by the action of creating two objects. In other words, the total number of ways that two objects do not change is given by the composition of the operators

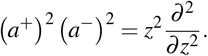

In order to understand the combinatorial interpretation, consider the action of (*a*^+^)^2^ (*a*^−^)^2^ on an arbitrary monomial such as *z*^5^, that is

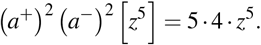

The combinatorial interpretation is: How many ways can we annihilate 2 objects out of 5 objects (*∂*^2^/*∂z*^2^) and then followed by bringing back 2 objects (*z*^2^). In other words, this action is nothing more than the permutation *P*(5, 2). Moreover, this means *P*(5,2) is the total number of ways that nothing has changed, hence this is how we calculate the number of ways the ‘Initial State” does not change.

Next, consider the action of (*a*^+^)^3^ (*a*^−^)^2^ on an arbitrary monomial such as *z*^7^, that is

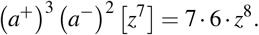

The combinatorial interpretation is: How many ways can we annihilate 2 objects out of 7 objects (*∂*^2^/*∂z*^2^) and then followed by bringing back 3 objects (*z*^3^). In other words, this action is how we calculate the “Final State.” The net change is defined as the Hamiltonian operator

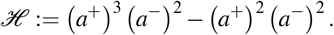

In general, the scenario where *j* distinct objects enter into an interaction and *k* objects emerge is shown in Figure 10. We describe these type of processes where the net flux is quantified by the change in the number of the objects via the Hamiltonian operator 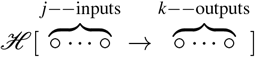. Towards this goal, the linear operator 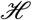 will be a modified Hamiltonian operator from quantum mechanics and will be composed of suitably modified creation and annihilation operators as has been defined in [2].

**Figure 10:**
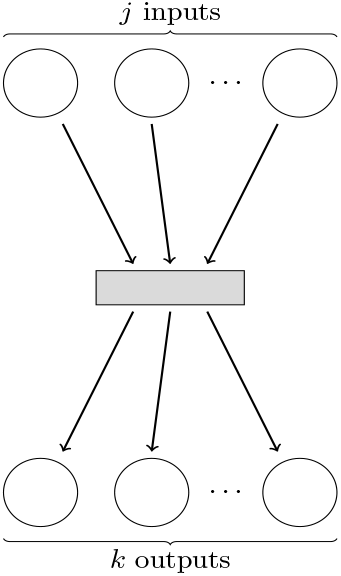
*j* Objects Enter into an Interaction & *k* Objects Emerge

Now that the Hamiltonian operator has been appropriately defined, the solution of an associated Schrödinger equation will describe how the probability of having exactly *n* objects at time *t* evolves over time.

First, consider the input side of the interactions. We quantify the scenario where all distinct input objects are annihilated followed by the action where all destroyed objects are then recreated. In other words, we are counting the total number of ways that the input configuration is unchanged. In general, if there are *j* objects in the initial configuration then the action is given by

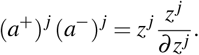

Next, examine the output configuration which is defined by annihilating *j* inputs and then recreating *k* outputs. The action of this process is given by

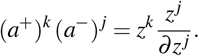

The stochastic Hamiltonian is defined in [2] as the difference between the final and initial configurations and is given by the stochastic Hamiltonian for the homogeneous class of *j*–inputs and *k*–outputs as follows:

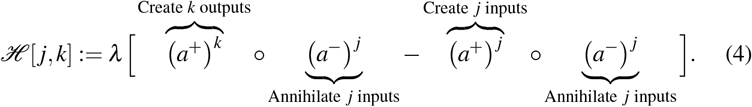

## 2 Quantum and stochastic mechanics

This section provides a short and self contained discussion of some of the tools of quantum/stochastic mechanics. The basic object describing the evolution in time of the probabilities of *n* distinct objects is the formal Markov generating function. Generating functions (**GF**) acts as a conduit between discrete and continuous mathematics.

> “A generating function is a clothesline on which we hang up a sequence of numbers for display.” [9]

The clothespins are the monomials *z^n^* and the individual laundry items are the probabilities *ϕ_n_*(*t*). A GF is written as an infinite power series where the coefficients of the monomials are the objects of interest. The monomials act as place holders and do not have any physical role in the analysis.

### 2.1 Generating functions

An ordinary^2^ generating function **GF** [9] is a formal power series of the form

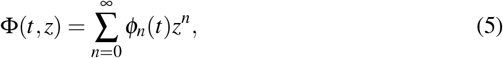

where the coefficients *ϕ_n_*(*t*) may or may not have physical meanings. Generating functions are extensively used in combinatorics, number theory, probability, and recurrence relations. The formal variable *z* has no physical meaning; it is basically a place holder. Additionally the analytic properties of the formal series **Φ**(*t, z*) will not be considered^3^.

In quantum mechanics the GF is defined as

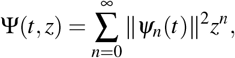

where the coefficients ||*ψ_n_*(*t*)||^2^ represent the amplitude of the wave function. In stochastic mechanics [2] the associated GF is defined in equation (5) where the density functions 0 ≤ *ϕ_n_*(*t*) ≤ 1 represent the probability of having exactly *n* objects at time *t*. Consider the special case where *z* = 1. In order to have a valid probability distribution, **Φ**(*t, z*) must satisfy the constraint

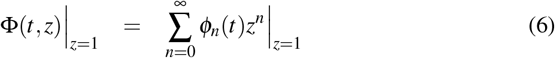

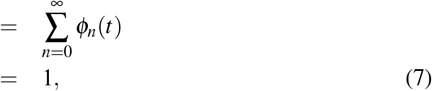

in which case

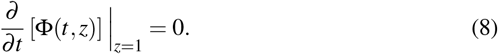

The master equation for the GF is given by

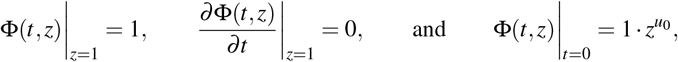

where 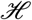 is the Hamiltonian operator associated with the interactions. The boundary and initial conditions are

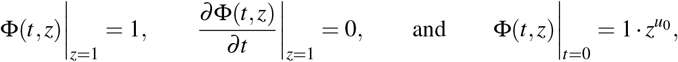

where 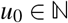 denotes the initial number of objects.

### 2.2 Expected value–first moment/mean of an observable

Recall that the expected value of a discrete probability distribution is given by

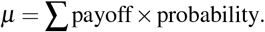

Suppose we are interested in the expected number of objects at time *t*. Consider the number operator defined as

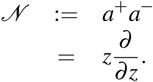

The reason it is called the number operator is that its action basically returns the number of objects as seen here

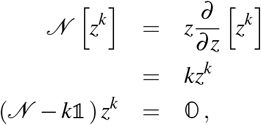

in which case

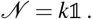

Now consider the action of the number operator 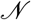 on the GF

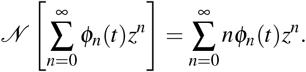

Next, define the expected number operator as

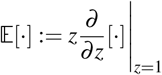

and lastly define the first moment *μ*(*t*) as

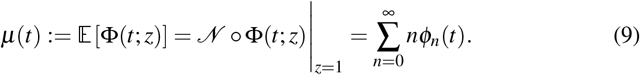

The standard macroscopic ODEs describing population dynamics will be derived and be of the form

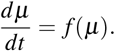

This quantum approach will surprisingly yield standard models such as the logistic equation.

### 2.3 Variance of an observable

The variance is defined as

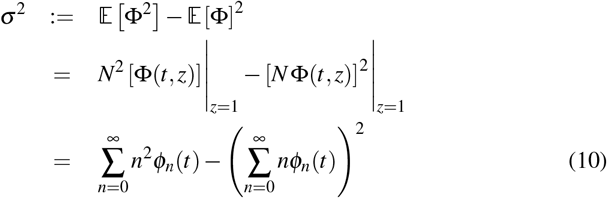

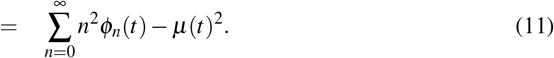

Another surprising result of this quantum approach is that the variance will also appear in the macroscopic ODEs. The implication is that noise, due to fluctuations in the interactions between members of the population, should also be included in the macroscopic model.

## 3 Immigration, natural death and fission

Consider a single species population undergoing the parallel processes of immigration, natural death, and fission as shown in Figure (11).

**Figure 11:**
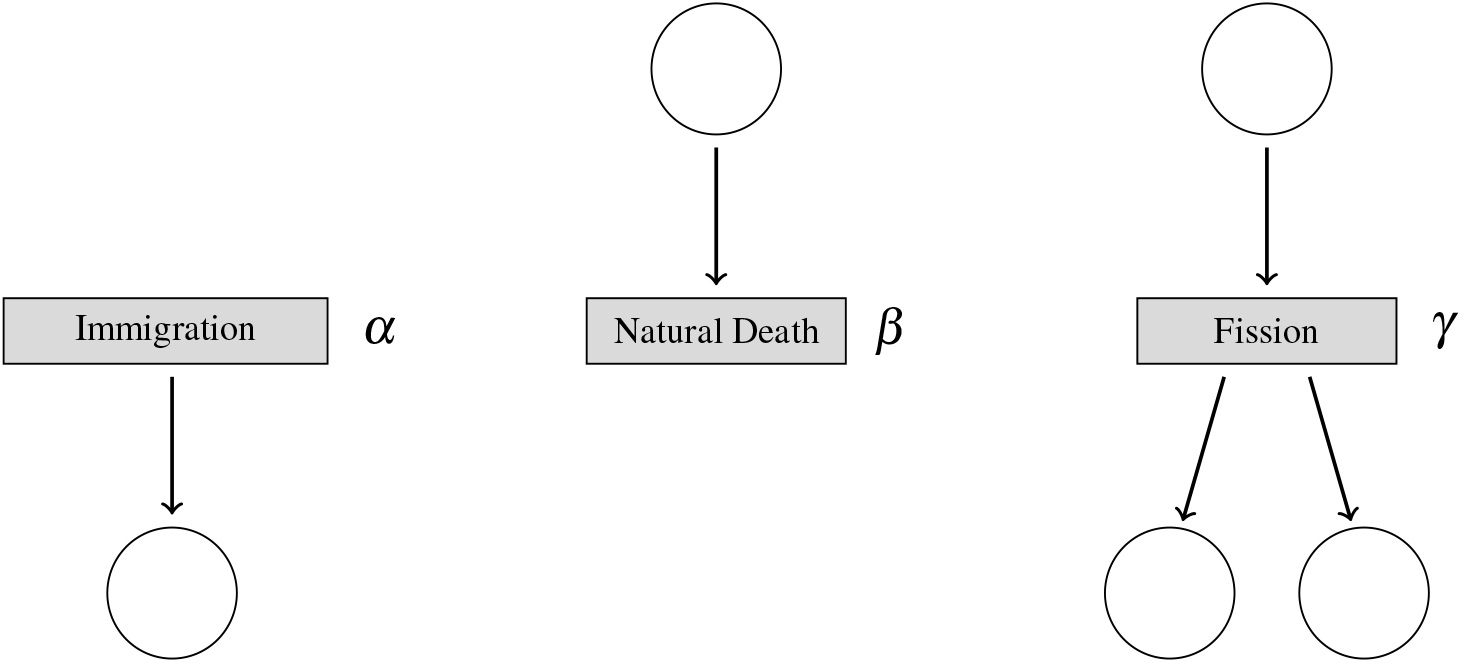
Immigration Birth (*α*), Natural Death (*β*), Fission (*δ*)

The parameters *α, β*, and *γ* are the rates (per unit time) at which each of these processes occur respectively. The immigration process has the associated Hamiltonian

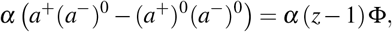

the death process operator is

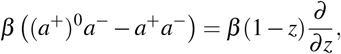

and the fission operator is

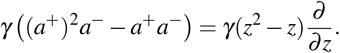

The associated master equation is given by

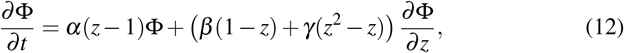

where **Φ**(*t, z*) is the GF of this Markov process.

### 3.1 Method of characteristics

In order to use the method of characteristics [10], rewrite the master equation as

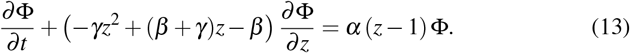

Assume that there exists differentiable parameterizations *t* = *t*(*r, s*) and *z* = *z*(*r, s*) such that

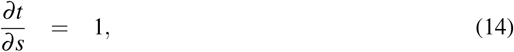

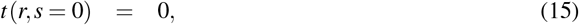

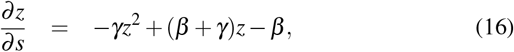

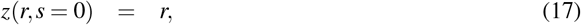

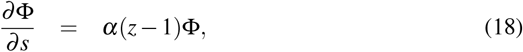

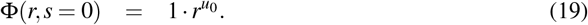

Integrating (14) and using the initial condition (15) yields *t* = *s*. Integrating (16) and using the initial condition (17) yields

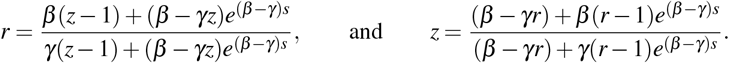

Lastly, integrating (18) and using the initial condition (19) yields the closed form ex-pression

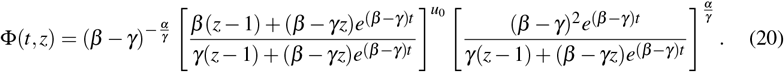

Using standard methods of analysis, it can be shown that the GF satisfies the essential properties

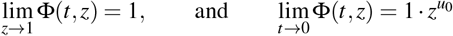

as well as the master equation given in equation (12).

### 3.2 Expected value

We now show that the quantum/stochastic paradigm predicts a familiar ODE model found in population dynamics. We find that *N***Φ** is

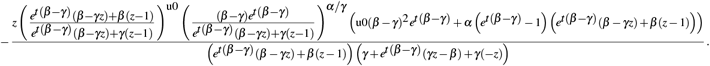

Evaluating at *z* = 1 yields the expected value is

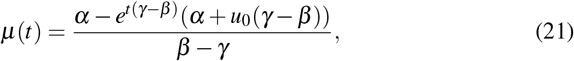

with initial condition *μ*(0) = *u*_0_. Notice that the first moment *μ*(*t*) satisfies the standard ODE population model

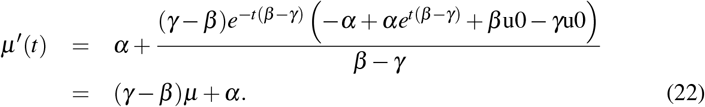

as found in single species population dynamics. The expression *γ* – *β* is a proxy for the net growth rate.

### 3.3 Second moment/variance of an observable

The variance is given by

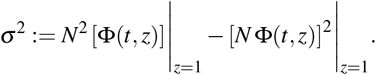

which is in fact

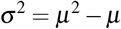

as expected.

### 3.4 Quantum tunneling of probabilities

We now discuss the implications of the quantum paradigm that cannot be deduced from the macroscopic viewpoint as given by the standard single species population model given in equation (22).

By expanding the explicit GF given in equation (20) the first two densities *ϕ*_1_(*t*) and *ϕ*_2_(*t*) are given by

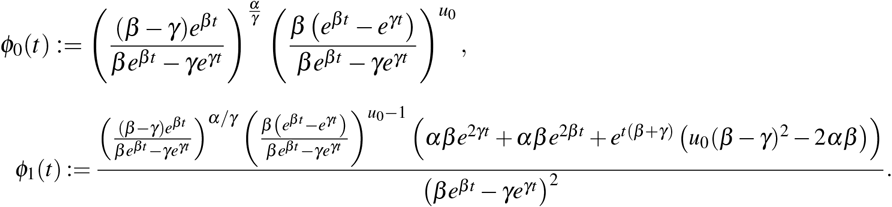

Obviously these discrete density functions are extremely complicated. In order to illustrate quantum tunneling of probabilities, we examine a special case. The above standard ODE model predicts that a stable equilibrium point is given by

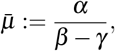

provided *γ* < *β*. If we choose *α* = *γ* = 1, *β* = 2, and *u*_0_ = 1 then the equilibrium point *μ* = 1 is stable and the GF reduces to the much simpler expression

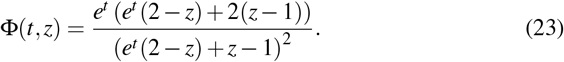

The individual densities reduce to

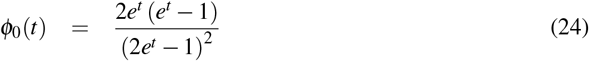

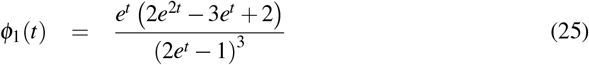

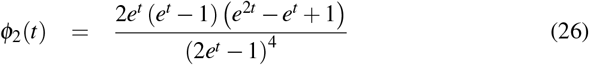

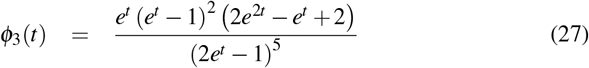

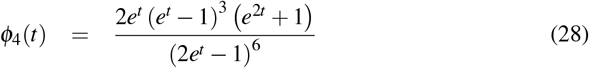

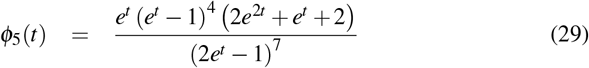

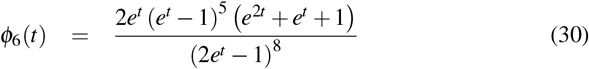

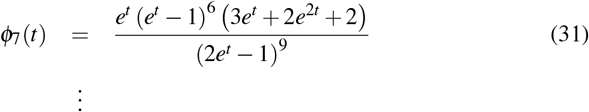

The graphs of the first 6 densities are shown in Figure (12). Note that the probabilities (as *t* → ∞) satisfy the decreasing monotonicity condition *ϕ*_*j*+1_ ≤ *ϕ_j_*.

**Figure 12:**
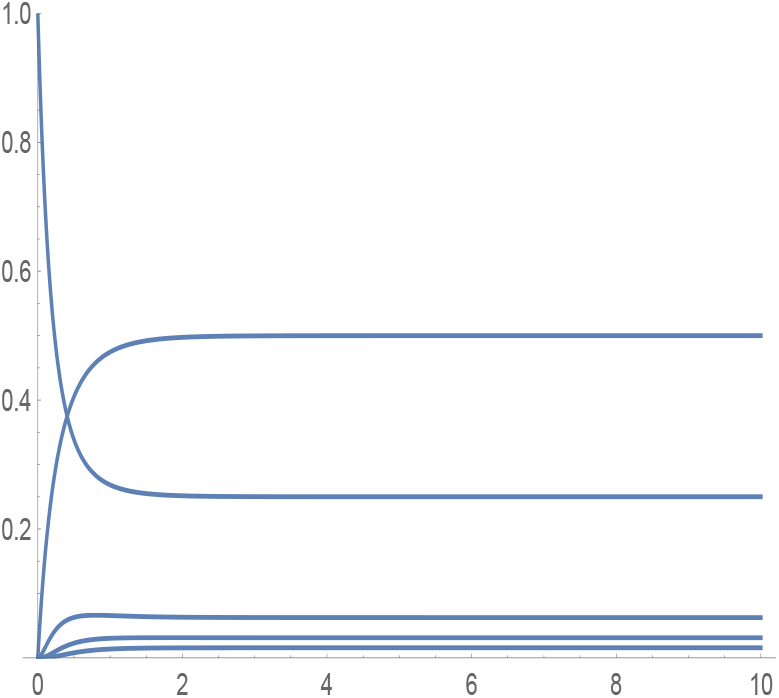
*ϕ*_0_,…, *ϕ*_5_

In fact, numerically it can be shown that 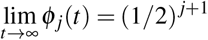 for *j* = 0,…, 17. Before rigorously proving this observation, we pause and compare the prediction made by the macroscopic model versus the quantum model. Specifically, if we start with one object, the macroscopic ODE model predicts that as *t* → ∞ the stable equilibrium point is 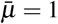. The quantum model however predicts that as *t* → ∞ the probability of having zero objects is 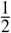, the probability of having one object is 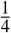. By extension, the probability of having 15 objects is 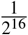, small but not zero. The quantum model given in equation (12) predicts a quantum tunneling effect of probabilities as a type of “noise” that is not captured by the standard deterministic ODE model given in equation (22).

### 3.5 Long–term behavior of the individual probabilities

In order to rigorously prove the above observation, substitute the GF into equation (12) with *α* = *γ* = 1, *β* = 2, and *u*_0_ = 1. Collecting the coefficients of the monomials *z^n^* yields the infinite system of first order ODE/difference equations

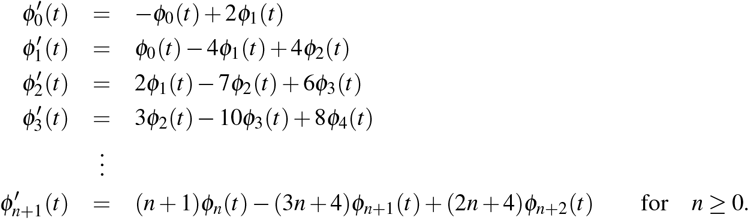

Since the GF can be written as 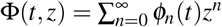 and the {*ϕ_n_*(*t*)} is a valid probability distribution, then 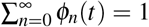 and so 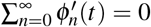 If a steady state exists for each density function then 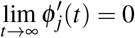. The infinite system of recurrence ODEs reduces to the infinite system of difference equations

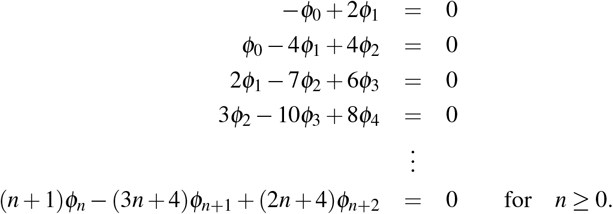

Using induction proves the desired result that 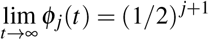 for 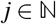.

## 4 ODE w/o closed form expression of Φ(*t, z*)

If the number of inputs *j* ≥ 2 then the associated PDE is of order *j* ≥ 2. In general, PDEs of 2nd order or higher cannot be solved explicitly for **Φ**(*t, z*). In this short section we show how to bypass having the explicit form of **Φ**.

The first moment is defined as 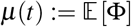, where

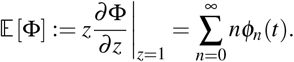

Differentiating *μ*

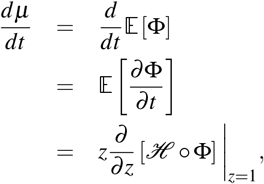

in which case

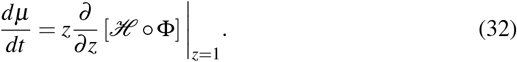

## 5 Immigration, death, competition and fission

We now derive a logistic ODE for the processes of immigration (rate *α*), natural death (rate *β*), competition (rate *γ*) and fission (rate *δ*). Since the Hamiltonian operator 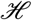 is a linear operator, we examine each of these processes individually and then add all the individual ODEs to obtain the initial value problem

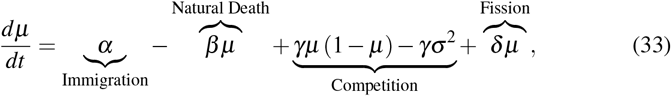

where *σ*^2^ denotes the variance.

### 5.1 Immigration

Consider the process of immigration with rate *α* as shown in Figure (13).

**Figure 13:**
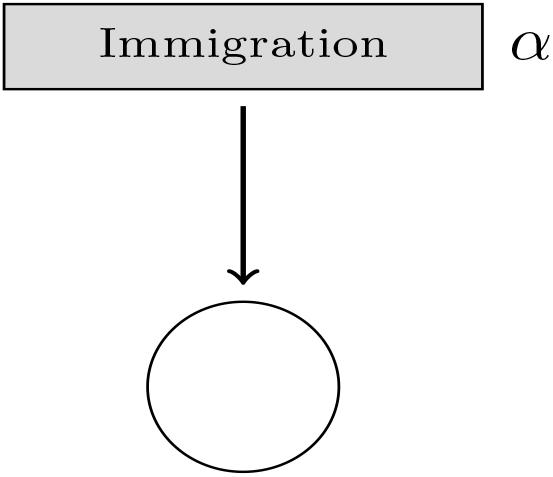
Immigration with rate *α*

The associated master equation is given by

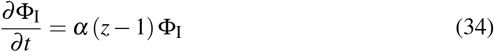

and using equation (32) we obtain the ODE

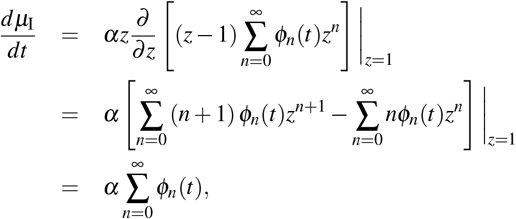

in which case

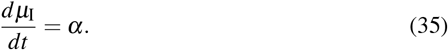

### 5.2 Natural death

Now consider the natural death process, as shown in Figure (14) with the associated master equation

**Figure 14:**
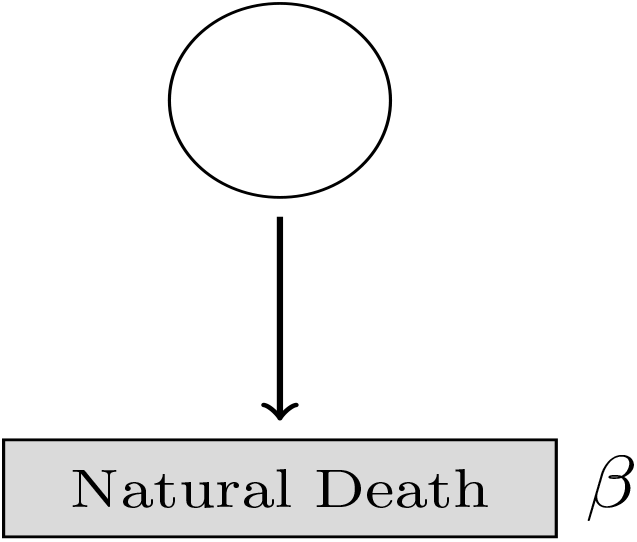
Natural Death With Rate *β*

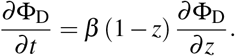

The associated macroscopic ODE is given by

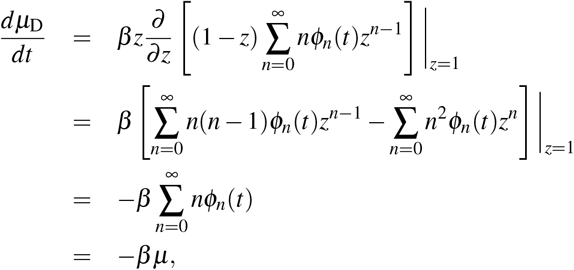

in which case

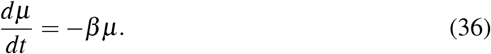

### 5.3 Competition

Next, consider the competition interaction process as shown in Figure (15) with the associated master equation

**Figure 15:**
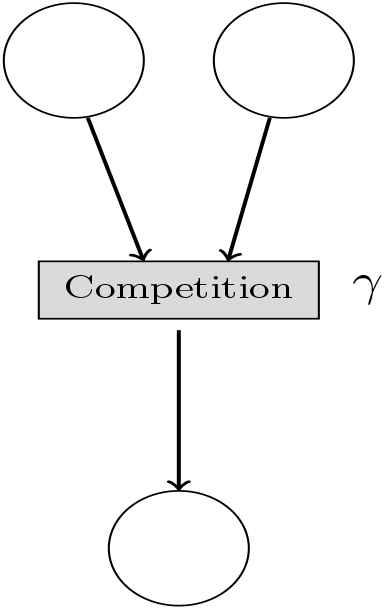
Competition With Rate *γ*

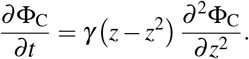

The macroscopic behavior is governed by the ODE

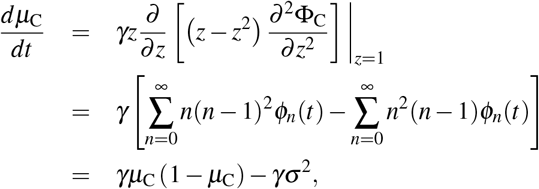

which yields the macroscopic description of competition

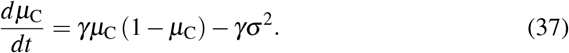

Consider the logistic equation with time dependent harvesting

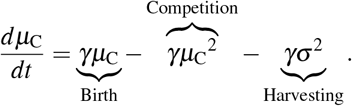

It is very interesting to note that the quantum formalism for a strictly decay process, namely competition, introduces two very unexpected terms in the macroscopic description. One would expect that there would not be any growth terms such as the intrinsic birth expression *γμ*_C_. Additionally, the quantum approach predicts a time dependent harvesting term via the variance expression *γσ*^2^. In other words, the quantum approach predicts that ad hoc birth and harvesting heuristic assumptions commonly included are actually justified by way of the effect of the quantum tunneling of probabilities.

### 5.4 Fission

Lastly, consider the process of fission as shown in Figure (16) Using the same methods as above yields the macroscopic description

**Figure 16:**
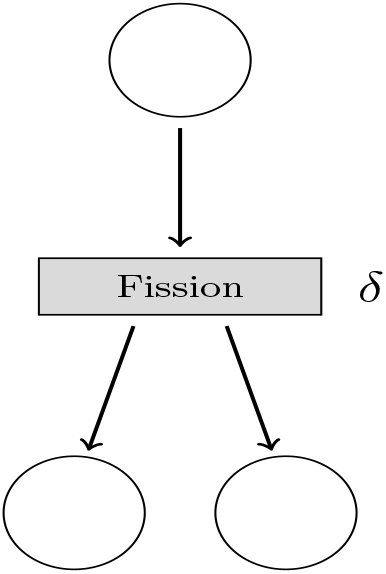
Fission With Rate *δ*

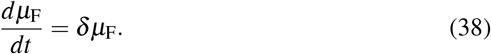

### 5.5 Equilibrium points & stability

Since each of the above Hamiltonian operators are linear, the macroscopic ODE is

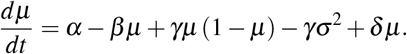

In order to discuss the equilibrium points as well as the stability, define 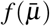, where

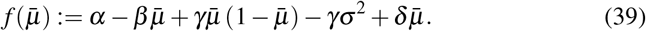

The positive equilibrium point

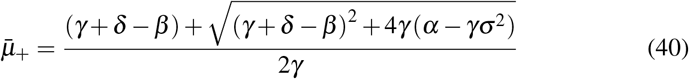

is ensured to be real and non-negative provided

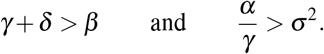

In other words, in order for the equilibrium point to be real and non–negative the rates must satisfy the constraints

Competition Rate+Fission Rate > Natural Death Rate and 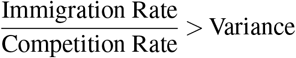.

Since

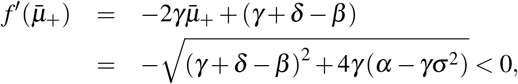

in which case 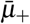 is stable. The one weakness in this analysis is that we do not have any knowledge on the temporal behavior of the variance *σ*^2^.

## 6 Concluding remarks

The quantum formalism, as defined in [2], is used to construct Schrödinger equations for single species population dynamics. The solution **Φ**(*t, z*) is given as a Markov generating function which describes the probability *ϕ_n_*(*t*) of having exactly *n* objects at time *t*. These probabilities exhibit quantum tunneling effects which predict events that are not seen or even expected in the standard deterministic models. The expected value of the solution **Φ**(*t, z*) yields similar deterministic models but with an additional noise term. This means that the quantum approach suggests that standard heuristic models lack one feature, namely noise. Furthermore, we have shown that the lone assumption of decay via competition results in the surprising result that a growth term occurs.

Future work will explore the use of these added features and may be helpful in predicting black-swan events. For example, consider a quantum model of HIV. Deterministic models, as they currently exist, do not allow the quantitative prediction of the possibility of a black-swan event such as an infected person actually surviving. This means that the quantum framework as discussed in this paper will need to be extended to multispecies interactions such as the standard SIR epidemiological model and the Lotka-Volterra predator-prey model. Additionally, we intend to explore the possibility of extending this framework to finite state automata with applications to gene regulatory networks. These mutations are eventually expressed as mutations. Lastly, including spatial aspects would yield spatial-temporal PDE models as well as the extension to cellular automata.

## Footnotes

1 For the physics community we use the symbol *a*^−^ instead of the commonly accepted notation *a*.

2 In this paper we will not discuss other generating functions such as Dirichlet, exponential, etc., gener-ating functions.

3 The reason for ignoring whether the series is/is not convergent is that the manipulations that will be performed are defined over the product topological ring of formal power series.

